# Decoding Prognostic Markers: Bioinformatics Insights into PIH1D1 and p53 in Children’s Brain Cancer

**DOI:** 10.1101/2023.12.09.570938

**Authors:** Dhiraj Kumar, Prashant Ranjan, Amit Kumar Verma, Riyaz Ahmad Mir

## Abstract

Genetic alterations in normal brain cells lead to the development of brain tumors (BT). The incidence of newly diagnosed cases is on the rise over time. Understanding the molecular biology of pediatric brain tumors is crucial for advancing novel therapeutic approaches to prevent or effectively manage this disease. The R2TP complex, a conserved co-chaperone from yeast to mammals, including RUVBL1, RUVBL2, PIH1D1, and RPAP3 in humans, plays a crucial role in the assembly and maturation of various multi-subunit complexes. This study evaluates the expression of PIH1D1 and p53 in pediatric brain cancers using The Cancer Genome Atlas (TCGA) data through the UALCAN platform—a novel, user-friendly resource for cancer OMICS data analysis.

Our analysis revealed elevated expression levels of PIH1D1 in pediatric brain tumors across all age groups compared to normal tissues, suggesting its potential as an early detection marker and a prognostic indicator. Additionally, P53 emerged as a promising target for brain tumor treatment, warranting exploration for age-specific applications. The Venn Diagram analysis further delineated key genes within the PIH1D1 and P53 networks, providing insights into the genetic landscape of childhood brain cancer.

Despite these findings, the study has limitations, primarily relying on bioinformatics databases. Future research will incorporate clinical tissues and cell lines in in vivo and in vitro experiments to unravel the mechanistic roles of PIH1D1 and p53 in brain tumors. This comprehensive approach bridges bioinformatics and clinical exploration, contributing to a holistic understanding of pediatric brain cancer molecular biology and paving the way for targeted therapeutic interventions.

## Introduction

Neoplasms that develop from various cell lineages are included in the wide variety of neoplasms that make up central nervous system tumours. Malignant gliomas and medulloblastomas (MB) are the most prevalent variations in the adult and paediatric populations, respectively. The most frequent solid tumour in children is a tumour of the central nervous system, which includes brain and spine tumours. The majority of cancer fatalities result from the 4,500 new brain tumours that are diagnosed each year. Low grade (less aggressive) or high grade (extremely aggressive) brain tumours are also possible. Primary brain tumours have no known aetiology, yet some of them contain germ line alterations and are frequently inherited. The majority are not inherited and are the product of somatic mutations(Philadelphia, 2014).

60 percent of paediatric tumours are found in the posterior fossa. The most frequent tumours are: diffuse intrinsic pontine glioma (DIPG), juvenile pilocytic astrocytoma (JPA), ependymoma, and atypical teratoid rhabdoid tumour (ATRT), in decreasing order of incidence. The cerebral hemispheres of the brain contain the other 40% of paediatric brain tumours. These include oligodendrogliomas, meningiomas, supratentorial primitive neuroectodermal tumours (PNET), astrocytomas, gangliogliomas, craniopharyngiomas, germ cell tumours, and dysembryoplastic neuroepithelial tumours (DNET)(Estevez-Ordonez et al., 2022, n.d.).

Gliomas, which are composed of the glial cells that make up the brain’s supporting tissue, are the most prevalent kind of brain tumour in people of all ages. Astrocytic and ependymal tumours are the two main subtypes of glial tumours. The most prevalent kind of juvenile glioma, astrocytomas, favour the neurological system. They generally take place in the cerebellum, a region of the brain that controls voluntary muscle movements and preserves equilibrium, posture, and balance. The majority can be treated surgically. In the optic nerve, astrocytes can develop, particularly in children with neurofibromatosis. Gliomas in the brainstem, located near the base of the brain, can potentially affect children. Ependymoma often develops from the cells lining the cerebral ventricles, which are cavities filled with cerebrospinal fluid. sluggish, frequently slow growing(Testa et al., 2018).

Identification of novel biomarkers in brain tumor is critical for accurate prognosis analysis and therapeutic efficacy prediction. In the past, a numbers of bioinformatics analyses have been conducted to identify key differentially expressed genes and enriched biological pathways or to evaluate the expression of a few specific genes in brain tumors, but such analysis using transcriptomes of brain tumours has not been satisfactorily performed. Therefore, it is urgently needed to discover novel molecular biomarkers, therapeutic targets, or prognostic evaluation index for brain tumour. The extensive application of bioinformatics databases has facilitated the discovery of new biomarkers for cancer management.

The structure of protein and RNP complexes appears to be a speciality of the R2TP complex, which is conserved from yeast to human. It participates in a variety of biological activities, including the production of small nucleolar ribonucleo proteins (snoRNPs), RNA polymerase, and PIKK signalling(Rivera-Calzada et al., 2017). It is made up of four distinct proteins, including PIH1D1/Pih1, RUVBL1/Rvb1, RUVBL2/Rvb2, and RPAP3/Tah1 (human/yeast). The AAA+ family of ATPases includes RUVBL1/Rvb1 and RUVBL2/Rvb2, and their physiologically relevant configuration seems to be an alternating hetero-hexamer/dodecamer. It is thought that PIH1D1/Pih1 and RPAP3/Tah1 heteromerize and serve as an adaptor to draw in clients as well as a link between HSP90 and the RUVBLs. The PAQosome is a structure that is made up of the R2TP and other prefoldin proteins in mammals(Houry et al., 2018). PIH1-N and PIH1-C are two of the domains covered by PIH1D1/Pih1. The DpSDD/E consensus sites are bound by the phospho-peptide binding PIH1-N region. Using proteomic and in silico screening, a variety of PIH1D1 phosphorylation-dependent binding partners were identified. One of them included the stability of the tumour suppressor p53 and was mediated by a motif related to the DpSDD sequence that has demonstrated direct phosphorylation-dependent interaction with the human protein ecdysoneless (ECD)(Hořejší et al., 2014). The protein p53 has received the most attention as a tumour suppressor. Apoptosis induction and cell cycle arrest are two of p53’s primary roles in the body. Additionally, p53 is involved in DNA repair, metabolic pathway regulation, embryo implantation, and cell senescence. Several viral oncoproteins inactivate the p53 gene. This opposes the idea that host cells undergo apoptosis to stop the spread and reproduction of viruses(Engeland, 2022).TP53 is the gene that is most frequently mutated in human malignancies. According to an estimate, this gene is mutated or deleted in almost half of tumours of all sorts on average. The bulk of other tumours are thought to have lost p53 activity through various mechanisms in addition to genetic inactivation. For instance, p53 proteolysis is brought on by viral oncoproteins or MDM2 overexpression.(Levine and Oren, 2009). Therefore, it is generally believed that the majority of tumours have lost p53 function either due to mutation or by the p53 pathway being compromised.

A transcription factor called p53 turns on a lot of genes. Both BAX and PUMA/BBC3 are prominent targets for its direct transcriptional activation. Importantly, almost half of the genes that p53 controls in terms of transcription are suppressed(Reisman et al., 2012). For a long time it was unresolved how p53 initiates downregulation of the many cell cycle regulators it controls, such as cyclin B1 and B2. It became evident that when we conducted genome-wide investigations in the setting of DREAM-dependent transcriptional repression, we discovered that p53 indirectly represses its many target genes. p21/WAF1/CIP1/CDKN1A is required for cyclin-dependent kinase inhibitor-mediated indirect transcriptional regulation by p53(Engeland, 2018). Despite the intensive study of PIH1D1 and p53 in many cancers, there is limited research regarding PIH1D1 in brain tumor. However, it remains also unclear how PIH1D1, a member of molecular chaperone complex contributes to oncogenesis. Currently, research groups and pharmaceutical companies are searching and developing inhibitors of R2TP/PAQosome as well as p53 as a promising chemotherapeutic target for cancers. However, the expression and prognostic values of PIH1D1 in brain tumor have not been well-studied. By studying the role of PIH1D1 and p53 in brain tumor, it might provide a novel biomarker for drug development and repositioning on this cancer.Herein, We had studied the expression patterns, interactive analysis, correlation expression in age group as well as in race.

## Methodology

### TCGA data analysis of PIH1D1 and p53 in Paediatric Brain cancer

The **U**niversity of **AL**abama at Birmingham **CAN**cer data analysis Portal (UALCAN) is a comprehensive, user-friendly, and interactive web resource for analyzing cancer OMICS data. It is built on PERL-CGI with high quality graphics using javascript and CSS. UALCAN is made to make it simple for users to access publicly available cancer OMICS data (TCGA, MET500, CPTAC, and CBTTC), as well as to identify biomarkers, validate potential genes of interest in silico, provide graphs and plots showing patient survival and expression profile, assess how promoter methylation regulates epigenetic regulation of gene expression, and perform pan-cancer gene expression analysis. Here we were evaluated the expression of PIH1D1 and p53 in paediatric brain cancer by TCGA analysis using data from The Children Brain Tumor Tissue Consortium (CBTTC) dataset. We had scan for PIH1D1 and p53 in dataset, then explored.

In this study Venny 2.1.0(Cheng et al., 2021) was used for Diagram analysis to identify common genes associated with childhood brain cancer. Firstly, a set of 12 common genes was determined by cross-referencing the PIH1D1 gene network obtained from GenMania(-GeneMANIA update 2018 | Nucleic Acids Research | Oxford Academic,? n.d.) with a refined gene list from GeneCards (Safran et al., 2010). This integrated approach facilitated the identification of a comprehensive roster of genes implicated in child brain cancer, providing a robust foundation for further investigation.

Simultaneously, a parallel analysis using the Venny 2.1.0 Diagram uncovered an additional 17 common genes associated with child brain cancer. The scrutiny of the P53 gene network sourced from GenMania played a central role in this process. The subsequent fine-tuning of gene selection was achieved through filtering with GeneCard, enhancing the specificity of the identified genes. The resultant compilation forms a key reservoir of candidates intricately tied to child brain cancer, benefitting from the shared attributes derived from both analytical sources.

This methodological approach leveraged the power of network analysis, cross-referencing, and filtering techniques to uncover genetic intersections in two distinct gene networks. The combination of these strategies aimed to provide a more comprehensive understanding of the molecular underpinnings of childhood brain cancer, setting the stage for further exploration and potential clinical implications.

## Results

Here We were evaluated the expression of PIH1D1 in pediatric brain cancer by TCGA analysis and observed that more or less similar pattern in both recurrence or progression as well as initial tumor (figure 1A) but p53 expression was low as compared to PIH1D1 in both case initial and recurrence(figure 1B). We analysed the expression of PIH1D1 was unchanged in both male and female children (figure 1C), whereas p53 (TP53) observed low in male and significantly high in female (figure 1D).

**Figure 1:**
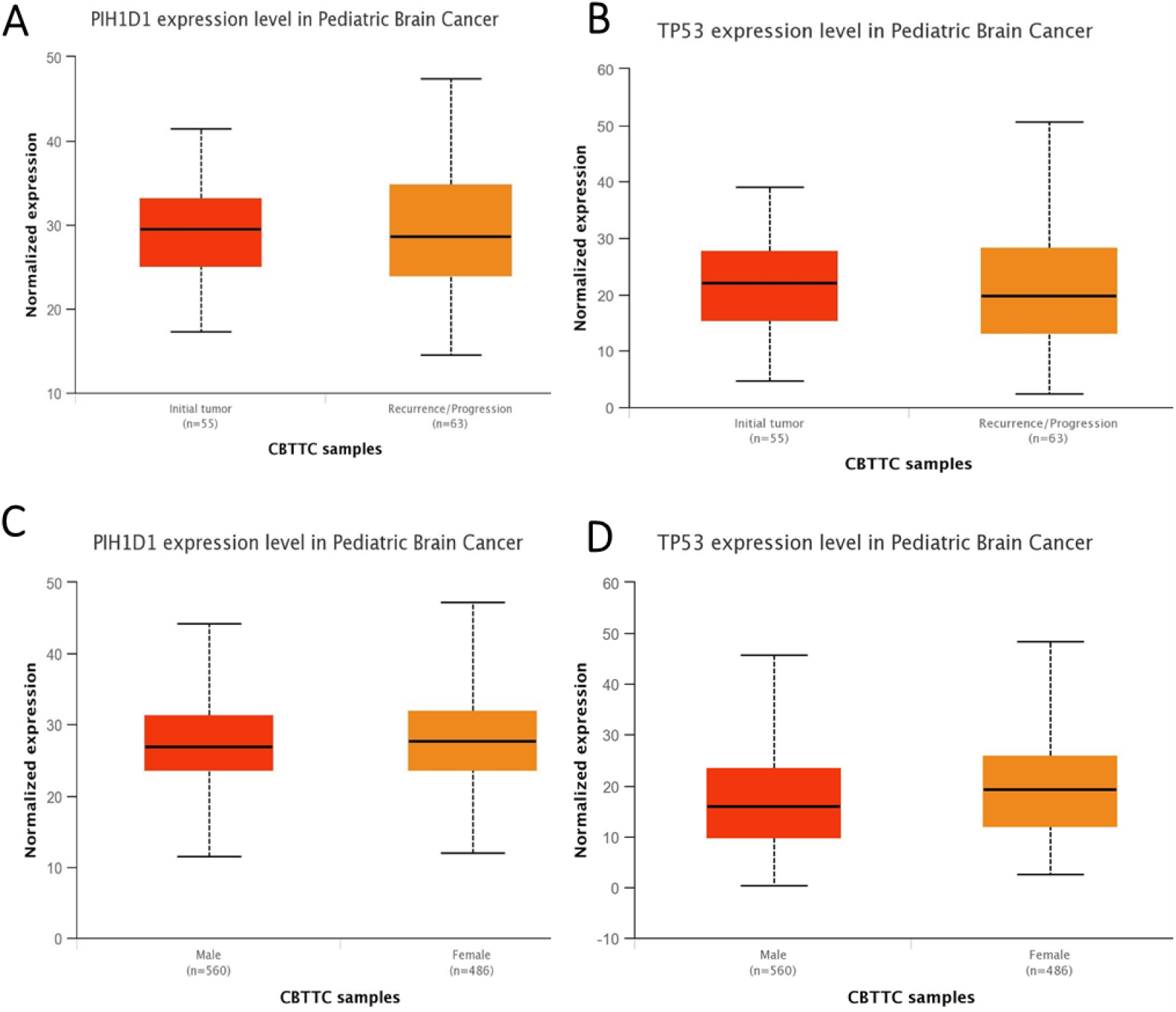
Expressional analysis of PIH1D1 and P53 in Pediatric brain cancer, A-B. PIH1D1 and P53 expression in initial tumour and recurrence, C-D. Gender wise expression of PIH1D1 and P53 in Pediatric brain cancer.

PIH1D1 expression was observed lowest in more than 30 year age group, whereas higher in up to 9 years age group (Figure 2A). Interestingly p53 found high in 20-29 years age group but Total expression level of p53 was low as compare to PIH1D1(Figure 2B). According to our analysis on basis of different races includes Caucasian, African-american, Asian, American Indian and other races, the PIH1D1 expression was observed approximately equal in all races whereas p53 expression was observed higher in Asian race but lower than PIH1D1 expression in Asian races(Figure 2 C&D).

**Figure 2:**
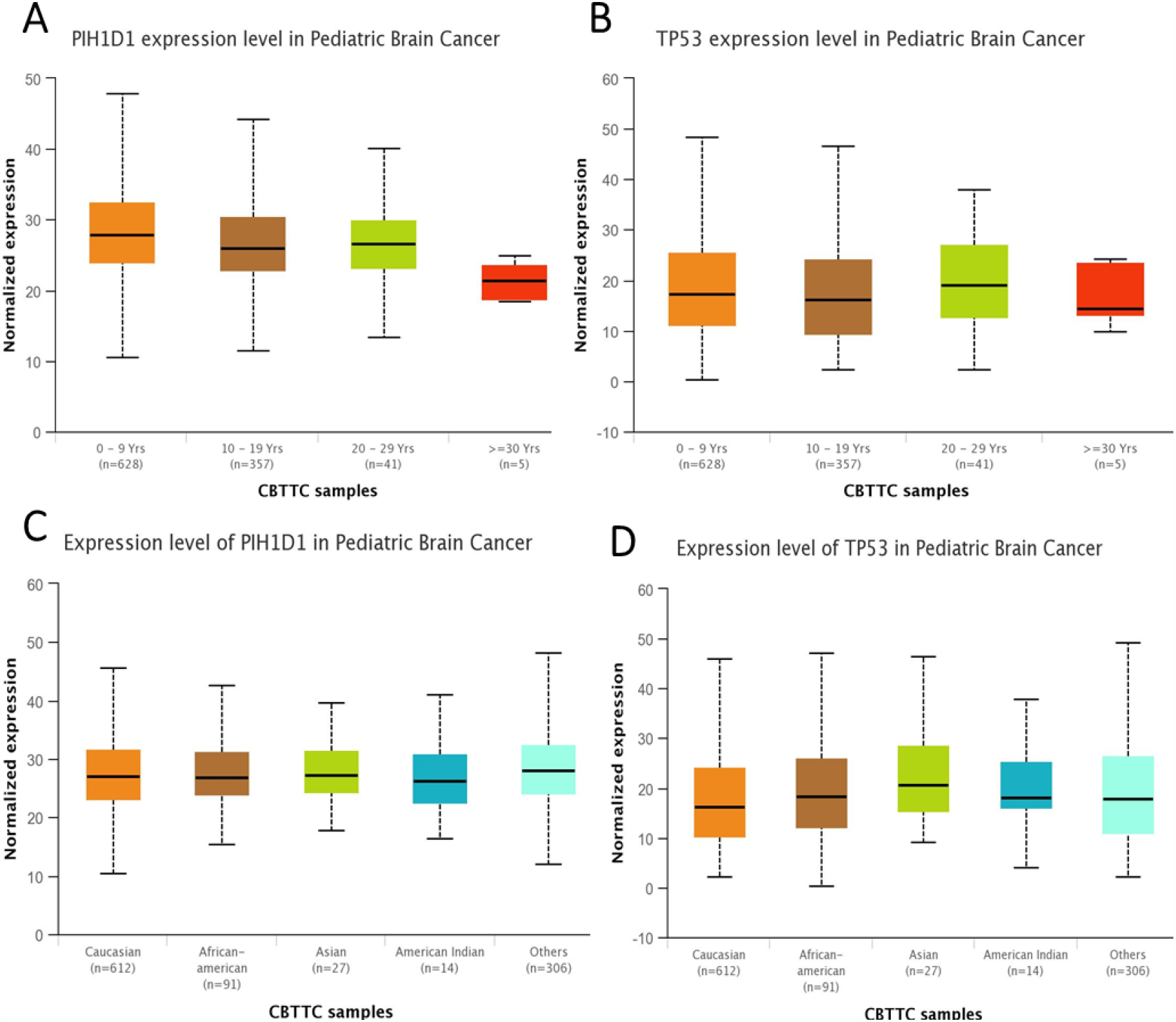
Expressional analysis of PIH1D1 and P53 in Pediatric brain cancer, A-B. Age wise expression of PIH1D1 and P53 in Pediatric brain cancer. C-D. Race wise expression of PIH1D1 and P53 in Pediatric brain cancer.

**Figure 3:**
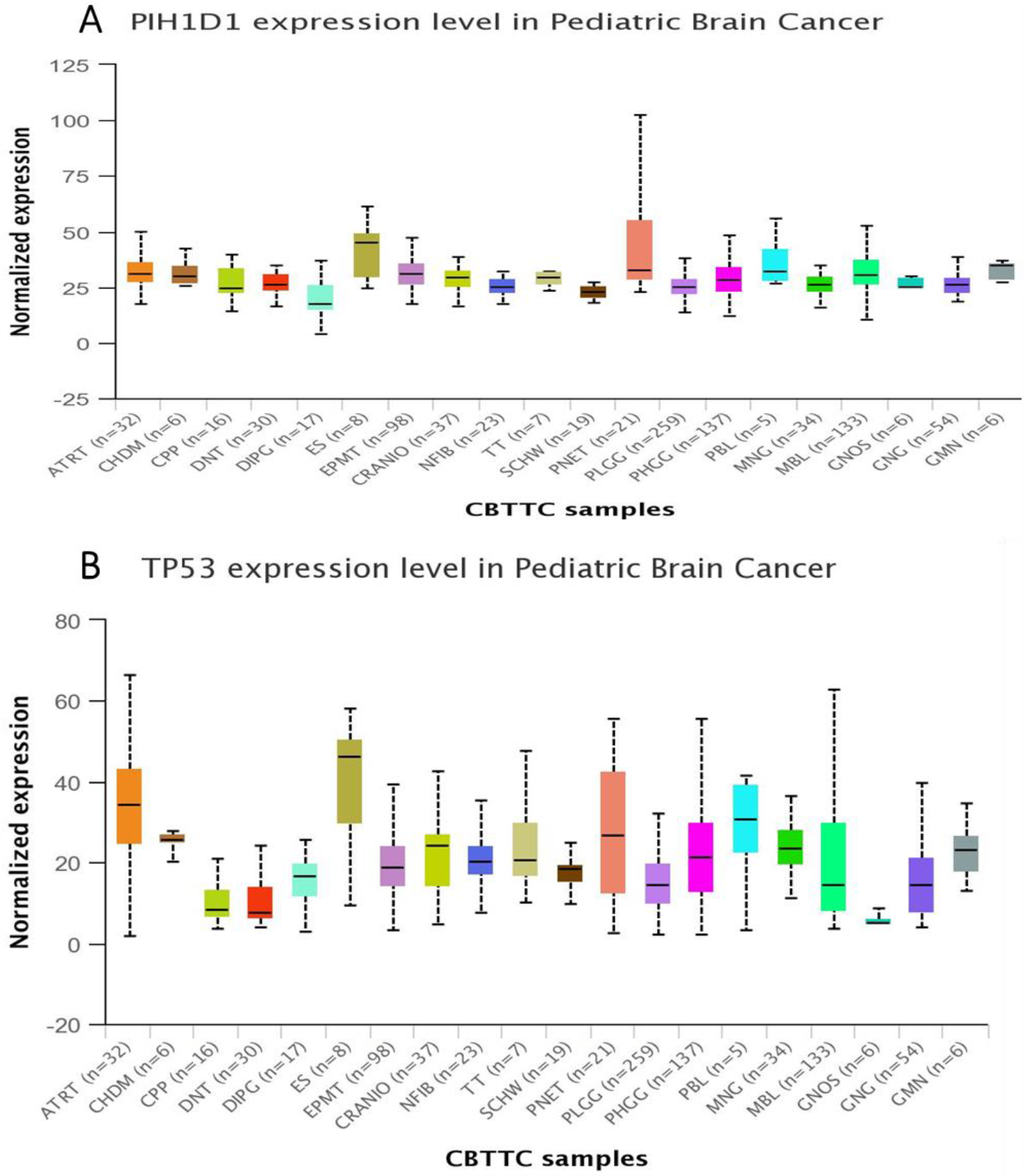
A. RNA expression profile of PIH1D1 in all the different type of Pediatric Brain cancer, B. RNA expression profile of P53 in all the different type of Pediatric Brain cancer.

**Figure 4:**
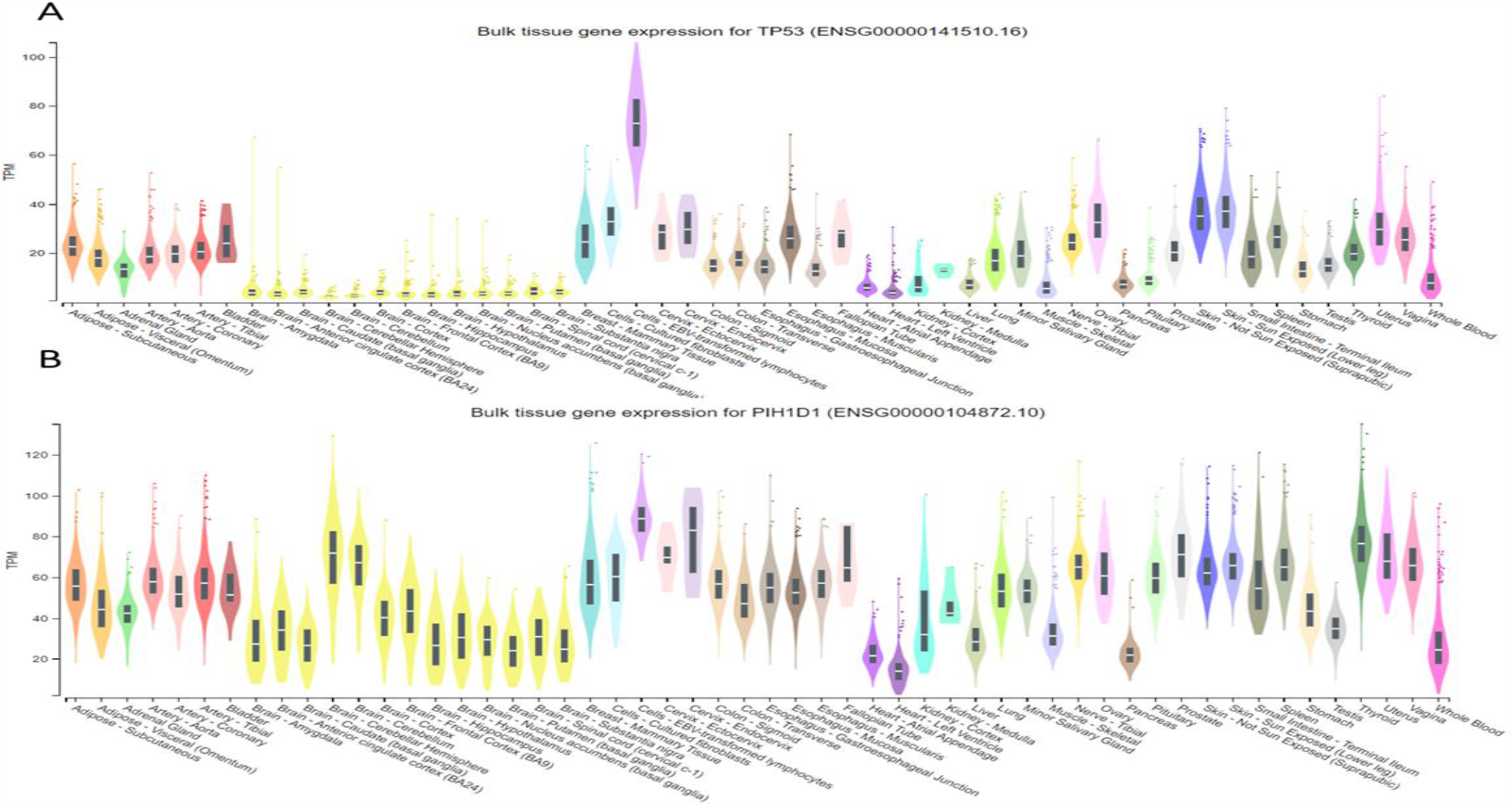
A. RNA expression profile of P53 in all the different type of Pediatric Brain cancer, B. RNA expression profile of PIH1D1 in all the different type of Pediatric Brain cancer.

RNA expression profile of PIH1D1 was analysed across all the different type of Brain cancer, and observed that Diffuse intrinsic pontine glioma (DPIG) shown to have lower expression and higher in Ewings Sarcoma (ES) and spinal cord primitive neuroectodermal (PNET) (Figure:3A), whereas p53 found lower expression in glial neuronal tumor not otherwise specified (GNOS) and choroid plexus papilloma (CPP) and higher in Ewings Sarcoma (ES), spinal cord primitive neuroectodermal (PNET) and Pineoblastoma(Figure:3B). All the abbreviation are given in Table 1.

**Table 1:** List of Abbreviated word.

| Abbreviation | Meaning | Abbreviation | Meaning |
| --- | --- | --- | --- |
| ATRT | Atypical Teratoid Rhabdoid Tumor | CRANIO | Craniopharyngioma |
| CPP | Choroid plexus papilloma | DNT | Dysembryoplastic neuroepithelial tumor |
| DIPG | Diffuse intrinsic pontine glioma | ES | Ewings Sarcoma |
| EPMT | Ependymoma | GNG | Ganglioglioma |
| GMN | Germinoma | MBL | Medulloblastoma |
| GNOS | Glial-neuronal tumor not otherwise specified (NOS) | NFIB | Neurofibroma/Plexiform |
| MNG | Meningioma | PHGG | High-grade glioma/astrocytoma (WHO grade III/IV) |
| PBL | Pineoblastoma | PNET | Supratentorial or Spinal Cord primitive neuroectodermal |
| PLGG | Low-grade glioma/astrocytoma (WHO grade I/II) | SCHW | Schwannoma |
| TT | Teratoma | CHDM | Chordoma |

Through Venn Diagram analysis, we pinpointed a collection of 12 common genes linked to childhood brain cancer. This set emerged by cross-referencing the PIH1D1 gene network acquired from GenMania with the gene list refined through GeneCards. The resulting intersection forms a comprehensive roster of pivotal genes implicated in child brain cancer (Fig 5&6).

**Figure 5:**
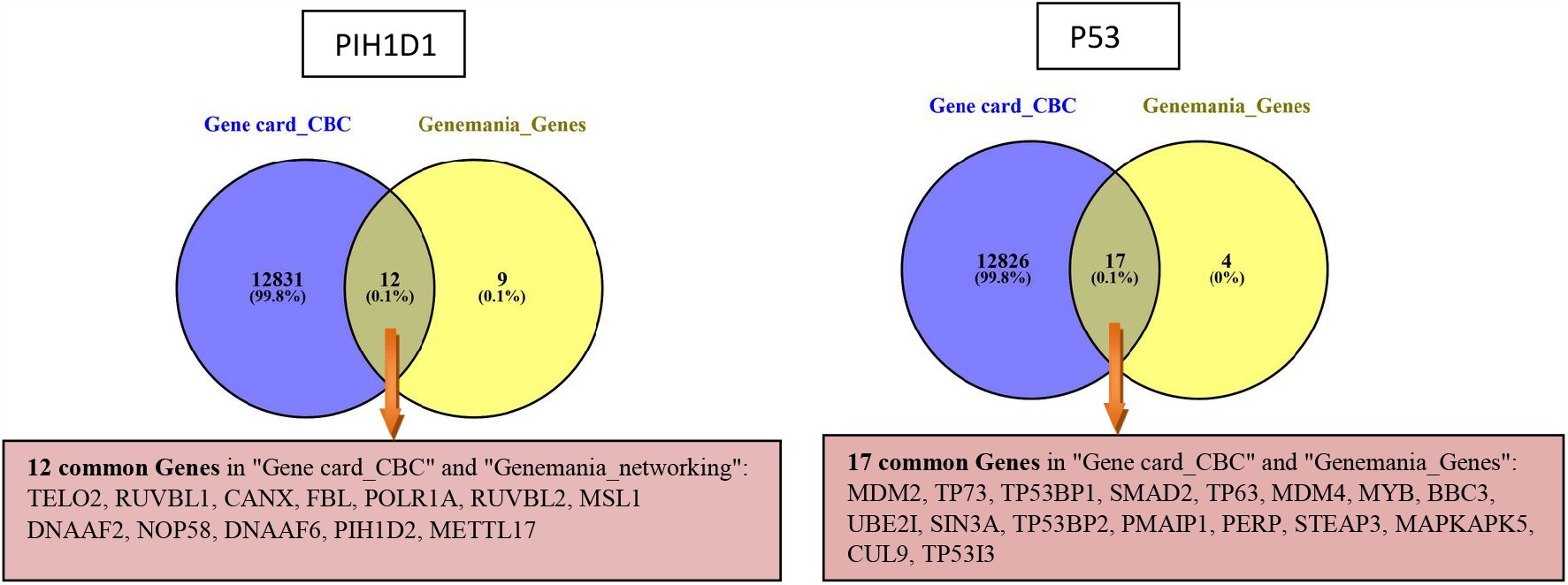
Harmonizing Networks: Deciphering 12 and 17 Key Genes in Child Brain Cancer through PIH1D1 and P53 Gene Network Analysis.

**Figure 6:**
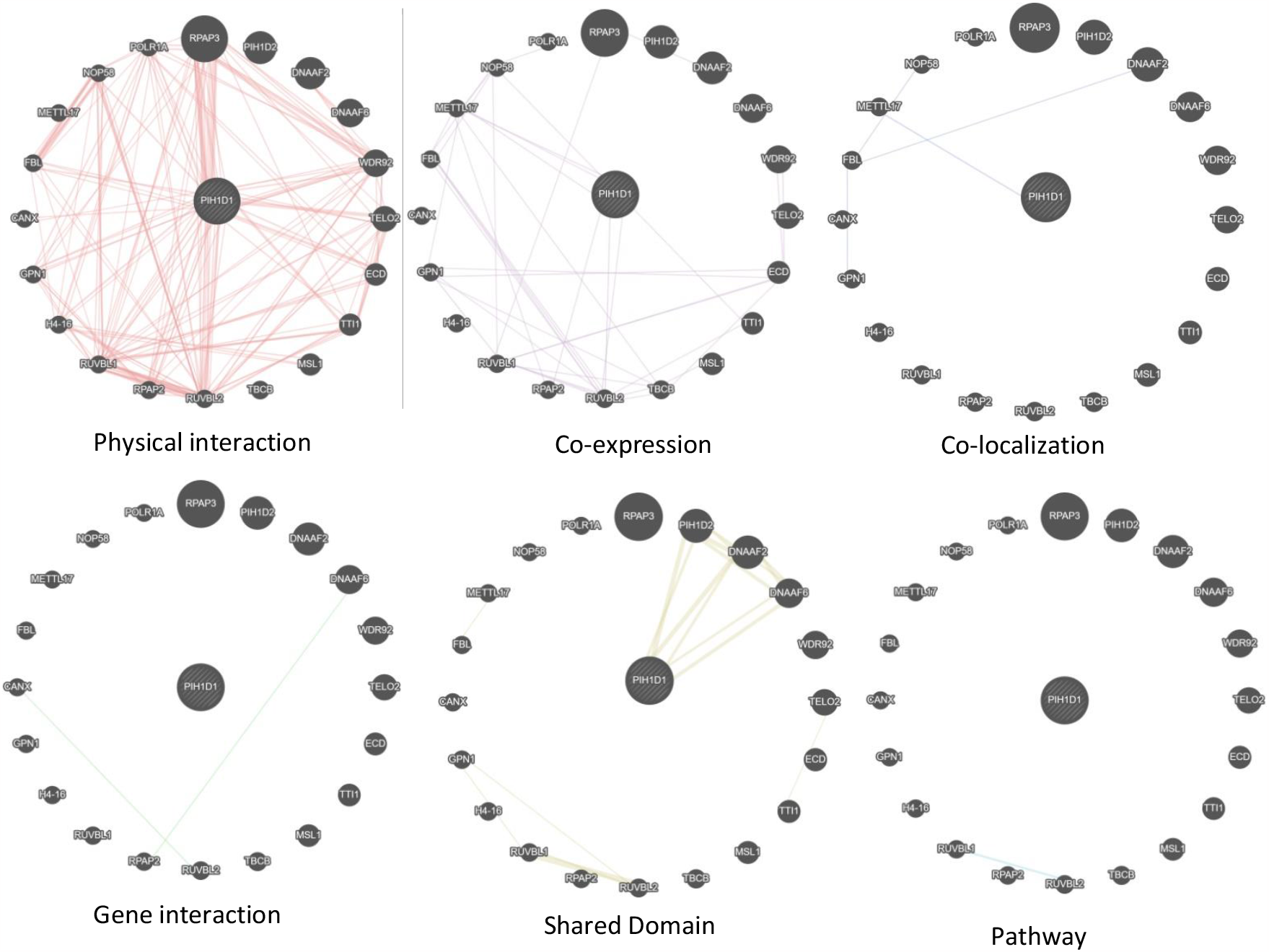
Bridging Networks: Unravelling Biomarker Potential through Interactive Analysis of PIH1D1

**Figure 7:**
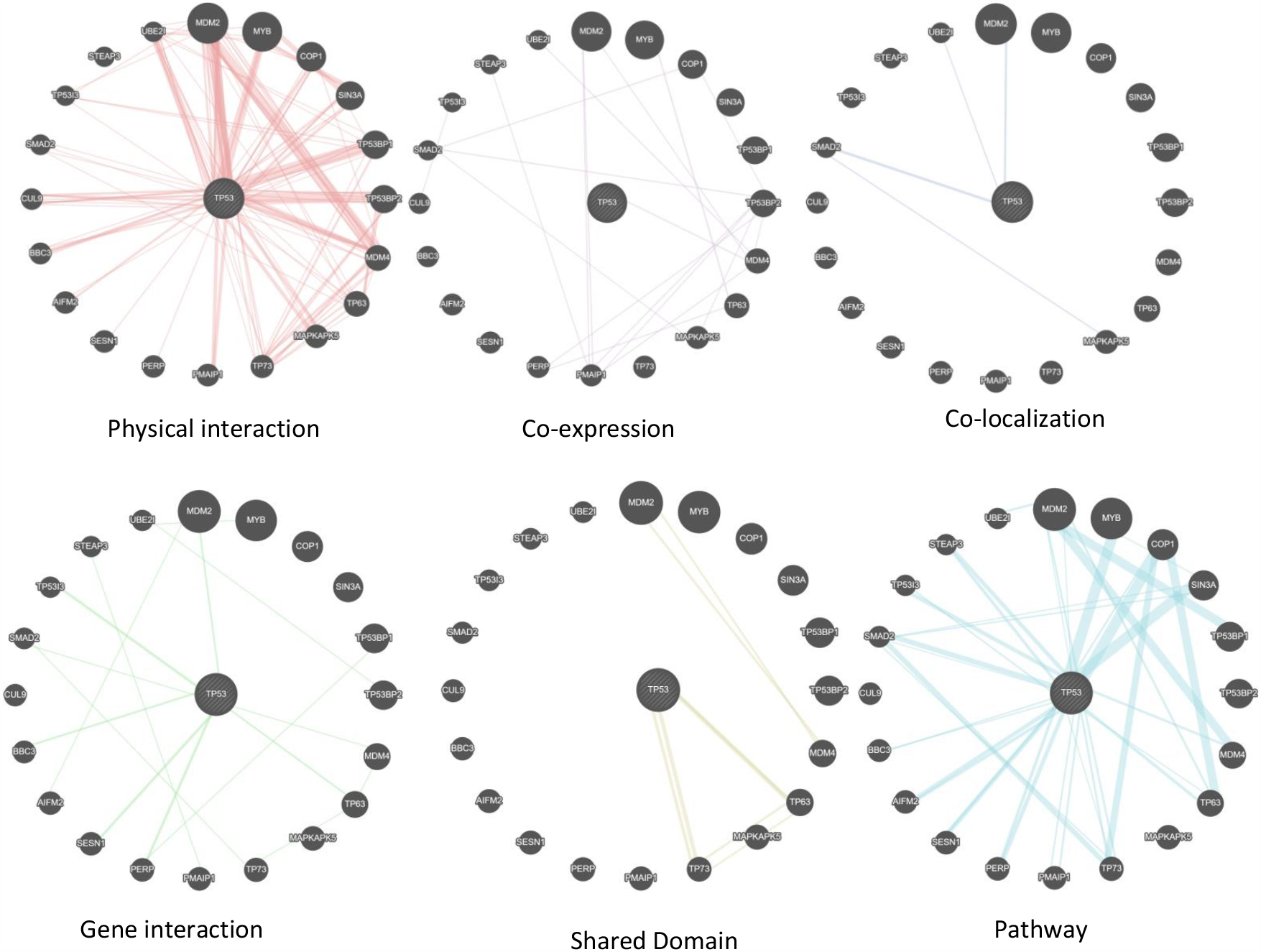
Network Nexus: Deciphering P53 Interactions for Biomarker Discovery

In a parallel analysis, the Venn Diagram revealed an additional 17 common genes associated with child brain cancer (Fig 5&7). These genes were discerned through scrutiny of the P53 gene network sourced from GenMania, and their selection was fine-tuned by filtering through GeneCard. The resultant compilation of genes serves as a key reservoir of candidates intricately tied to child brain cancer, drawing on shared attributes from both analytical sources.

## Discussion

The most prevalent solid tumours in children are malignant brain tumours. They account for 20–30% of all children malignancies and are the main cancer-related killer in this age range.(Baldwin and Preston-Martin, 2004)The most often mutated gene in human cancer is the p53 tumour suppressor gene (TP53), which is also present in different forms of brain tumours(Fulci et al., 2006). One of the key contributing factors to the emergence of cancer is p53 function loss. It has been established that PIH1D1 has a role in the control of apoptosis(Inoue et al., 2010).The protein PIH1D1 is known to be unstable, and In a previous study it was shown that Tah1 (RPAP3 in humans) is another R2TP complex component that stabilises PIH1D1(Kamano et al., 2013). Although the role of PIH1D1 in the tumorigenesis and prognosis of several cancers has been partially confirmed, further bioinformatics analysis of Brain tumor has yet to be performed. Importantly, the current study focuses on the analysis and exploring the expression and prognostic values of PIH1D1 and p53 in Brain tumor.

In our study, we found that the expression of P53 was significantly higher in Brain tumor than PIH1D1 in both case initial and recurrence.Similarly we analysed the expression of PIH1D1 was unchanged in both male and female children, whereas p53 (TP53) observed low in male and significantly high in female.We also observed that both PIH1D1 and p53 expression were low in more than 30 year age group, whereas higher in all the age group. If we focus on up to 9 years age group, we observed PIH1D1 expression was higher. Interestingly p53 found higher in 20-29 years age group but Total expression level of p53 was low as compare to PIH1D1. These data suggest PIH1D1 can be used for early detection of Brain tumor in peads and may be used as indicators of prognosis. Moreover, P53 can be used as the target of younger or later age for Brain tumor in the future.

According to our analysis on basis of different races includes Caucasian, African-american, Asian, American Indian and other races, we observed no difference in the expression of PIH1D1 whereas p53 expression was observed higher in Asian race.PIH1D1 mRNA expression profile was analysed across all the different type of Brain cancer, and observed that Diffuse intrinsic pontine glioma (DPIG) shown to have lower expression and higher in Ewings Sarcoma (ES) and spinal cord primitive neuroectodermal (PNET), similarly p53 also found higher expression in Ewings Sarcoma, and lower expression in glial neuronal tumor not otherwise specified (GNOS) and choroid plexus papilloma (CPP).We compared the expression levels of PIH1D1 mRNA across cancers and its mRNA expression with that in corresponding normal tissues and found that PIH1D1 expression was higher in almost all cancer groups than in normal tissues, including the brain, breast, and head and neck however lower in Kidney Renal Clear Cell Carcinoma (KIRC). Whereas the mRNA expression of p53 was significantly downregulated in all different types of brain tumours. In previous study it was observed in both clinical samples and the TCGA database, the expression of several additional genes, including NR1B2 in KIRC, was markedly down-regulated(Yin et al., 2019).

The Venn Diagram analysis conducted in this study has provided valuable insights into the genetic landscape of childhood brain cancer, revealing distinct sets of common genes associated with the disease through the PIH1D1 and P53 gene networks.

The identification of 12 common genes associated with childhood brain cancer, achieved by cross-referencing the PIH1D1 gene network from GenMania with the gene list refined through GeneCards, underscores the importance of collaborative data integration. This approach has allowed us to pinpoint a comprehensive roster of genes that may play pivotal roles in the onset and progression of child brain cancer. The synergy between GenMania and GeneCards in this analysis enhances the robustness of our findings, providing a foundation for further investigation into the specific functions and interactions of these genes within the context of childhood brain cancer.

In a parallel exploration, the Venn Diagram unveiled an additional 17 common genes associated with child brain cancer, identified through the scrutiny of the P53 gene network from GenMania. The meticulous fine-tuning of gene selection through filtering with GeneCard adds a layer of specificity to our results. The compilation of genes derived from this analysis forms a reservoir of candidates intricately linked to child brain cancer. The shared attributes drawn from both the PIH1D1 and P53 networks suggest a potential convergence of molecular pathways contributing to the disease, highlighting the complexity of its genetic underpinnings.

The amalgamation of findings from both analyses enriches our understanding of the genetic factors associated with childhood brain cancer. The shared genes identified in both sets may serve as focal points for future research, warranting further investigation into their functional significance and potential as diagnostic or therapeutic targets. It is essential to acknowledge the limitations of this study, including the need for experimental validation and consideration of the heterogeneous nature of childhood brain cancer subtypes.

## Conclusion

In summary, this analysis shows that both PIH1D1 and p53 expression were low in more than 30 year age group, whereas higher in all the age group. The expression of PIH1D1 was unchanged in both male and female children, whereas p53 (TP53) observed low in male and significantly high in female. We suggest PIH1D1 can be used for early detection of Brain tumor in peads and may be used as indicators of prognosis. Moreover, P53 can be used as the target of younger or later age for Brain tumor in the future.

## Acknowledgments

D.K. is thankful to AIIMS New Delhi-110029 and Jamia Millia Islamia, New Delhi 110025 for providing necessary facilities to carry out this piece of work.

## Data Availability

Mainly Data of Brain tumor was downloaded for this study from TCGA website, UALCAN datasets.

